# The immunosuppressive Tuberculosis-associated microenvironment inhibits viral replication and promotes HIV-1 latency in CD4+ T cells

**DOI:** 10.1101/2023.12.05.570223

**Authors:** Samantha Cronin, Anneke de Vries-Egan, Zoï Vahlas, Alejandro Czernikier, Claudia Melucci, Pehuén Pereyra Gerber, Thomas O’Neil, Brian Gloss, Mayssa Sharabas, Gabriela Turk, Christel Verollet, Luciana Balboa, Sarah Palmer, Gabriel Duette

**Affiliations:** The Westmead Institute for Medical Research, Centre for Virus Research, Westmead, Australia; University of Sydney, Faculty of Medicine and Health, Sydney Australia; Institut de Pharmacologie et Biologie Structurale (IPBS), Université de Toulouse, Centre National de la Recherche Scientifique, Université Toulouse III - Paul Sabatier (UPS), Toulouse, France; International Research Project CNRS “ MAC-TB/HIV ”, Toulouse, France and Buenos Aires, Argentina; Instituto de Investigaciones Biomédicas en Retrovirus y SIDA, Universidad de Buenos Aires-CONICET, Buenos Aires, Argentina; Cambridge Institute for Therapeutic Immunology and Infectious Disease, Jeffrey Cheah Biomedical Centre, Cambridge, United Kingdom; Instituto de Medicina Experimental-CONICET, Academia Nacional de Medicina, Buenos Aires, Argentina

## Abstract

*Mycobacterium tuberculosis* (*Mtb*), the causative agent of tuberculosis (TB), is the most common coinfection among people living with HIV-1. This coinfection alters the efficacy of the immune response against both HIV-1 and *Mtb*, and is associated with accelerated HIV-1 disease progression and reduced survival. Enhanced HIV-1 replication in macrophages induced by *Mtb* coinfection may contribute to the worsened clinical outcomes observed in HIV-1/TB coinfected individuals. However, the impact of the HIV-1/TB coinfection on HIV-1 replication and latency in CD4+ T cells remains poorly studied.

In this study, we used the acellular fraction of tuberculous pleural effusion (TB-PE) as a proxy for the microenvironment generated by *Mtb* infection. Using this physiologically relevant fluid, we investigated whether viral replication and HIV-1 latency in CD4+ T cells are affected by a TB-associated microenvironment. Interestingly, our results revealed that TB-PE shaped the transcriptional profile of CD4+ T cells impairing T cell receptor-dependent cell activation and decreased HIV-1 replication. Moreover, this immunosuppressive TB microenvironment promoted viral latency and inhibited HIV-1 reactivation in CD4+ T cells from people living with HIV-1. This study indicates that the immune response induced by TB may contribute to the persistence of the viral reservoir by silencing HIV-1 expression in individuals coinfected with both pathogens, allowing the virus to persist undetected by the immune system and increasing the size of the HIV-1 latent reservoir in cells at the site of the coinfection.

## Introduction

Despite decades of research and the effectiveness of antiretroviral therapy (ART), HIV-1 remains an incurable infection. Globally in 2022, there were more than 38 million people living with HIV-1 (PLWH), with 630,000 individuals dying from HIV-1-related causes (1). The major obstacle to an HIV-1 cure is the latent HIV-1 reservoir, as replication-competent HIV-1 proviruses persist within long-lived memory CD4+ T cells of ART-suppressed individuals (2–6). These latent proviruses are largely transcriptionally silent and hence remain hidden from immune surveillance and CD8+ T cell-mediated clearance. However, the virus can rebound from these latently HIV-infected cells when ART is interrupted (7–10).

A potential curative strategy for HIV-1 is called ‘shock and kill’, which involves reactivation of HIV-1 from latently infected cells (shock), and subsequent elimination (killing) of these cells, through the combined effects of HIV-1-specific cytotoxic CD8+ T cells and ART (11). Therefore, the success of these curative strategies in clearing HIV-1 from PLWH relies on the ability to broadly reactivate HIV-1 from latently infected cells.

*Mycobacterium tuberculosis* (*Mtb*), the causative agent of tuberculosis (TB), ranks alongside HIV-1 as a leading infectious killer. Over 10 million new cases of TB were identified in 2021, with a total of 1.6 million deaths attributed to TB infection (12). In addition, an estimated one-third of HIV-1-infected individuals are coinfected with *Mtb*, with PLWH being 18 times more likely to develop active TB disease than HIV-1-negative individuals (12). Importantly, PLWH and TB experience accelerated HIV-1 disease progression and reduced survival in comparison to people living with HIV-1 alone (12, 13). As such, TB remains the leading cause of death amongst PLWH, with 187,000 deaths attributed to HIV-1/TB coinfection in 2021 – accounting for one in three Acquired Immunodeficiency Syndrome (AIDS)-related deaths (12, 14).

It has been previously shown that infection with *Mycobacterium tuberculosis* (*Mtb*) or mycobacterial component(s) may promote HIV-1 replication (15–21). Moreover, it has recently been demonstrated that TB-associated microenvironments, such as therapeutically aspirated pleural effusions (PEs) from TB patients (TB-PE), render human macrophages highly susceptible to *Mtb* infection (22, 23), as well as to HIV-1 infection and spread (21, 24). However, contradictory evidence has also emerged showing inhibition of HIV-1 replication by *Mtb*, blurring our understanding of the interaction between these two pathogens (25, 26). Furthermore, whether *Mtb* influences viral replication and/or HIV-1 latency, specifically in CD4+ T cells, remains unclear.

Pleural effusion is an excess of fluid recovered from the pleural space of the human respiratory compartment and is found in approximately 30% of TB patients (27). This excess of fluid is caused by the spread of *Mtb* into the pleural space, leading to subsequent local inflammation and leukocyte infiltration (28). Importantly, in HIV-1/TB coinfected persons, the formation of PE is more common and contains high HIV-1 titers, compared to serum from the same individual (29–31). Thus, TB-PE is a physiologically relevant fluid that can be used as an *ex vivo* model for a human compartment impacted by *Mtb* infection (21, 22, 24, 32). Therefore, in this study we investigate the effects of TB-PE on viral replication and HIV-1 latency in CD4+ T cells.

## Results

### Exposure to TB pleural effusion inhibits HIV-1 replication in primary CD4+ T cells

Whether the inflammatory microenvironment generated by the host immune response to *Mtb* infection impacts HIV-1 replication in CD4+ T cells remains undetermined. To address this question, primary CD4+ T cells were incubated with 20% v/v TB-PE (pooled from 9 participants with TB and devoid of viable mycobacteria) for 1-2 hours and then infected with HIV-1. The proportion of infected cells was determined by HIV-1-p24 staining and flow cytometry at day 3 post-infection (Figure 1A). Of note, we did not observe differences in cell viability across the different treatments (Figure 1B). We found that the pre-treatment with TB-PE significantly decreased the proportion of HIV-1 infected cells in primary CD4+ T cells, as shown in Figure 1C. Importantly, this result was reproduced when cells were treated with TB-PE collected from two individual TB patients (Figure 1D). In contrast, pre-treatment with PE from individuals who experienced heart failure (HF-PE) did not exhibit a significant inhibition of HIV-1 infection, indicating that the inhibition of HIV-1 replication is dependent on the PE etiology (Figure 1E). To assess whether the effect of TB-PE on HIV-1 infection is reversible, we washed the TB-PE-treated infected cells 24 hours after infection, and replaced the media with fresh media devoid of TB-PE. At day three post infection, we observed an increase in the number of p24-positive cells in the cultures where TB-PE was removed, as compared to cells that remained exposed to TB-PE (Figure 1F), indicating that the inhibition of HIV-1 infection induced by TB-PE is reversible. Our results show that the TB microenvironment decreases HIV-1 replication in CD4+ T cells.

**Figure 1.**
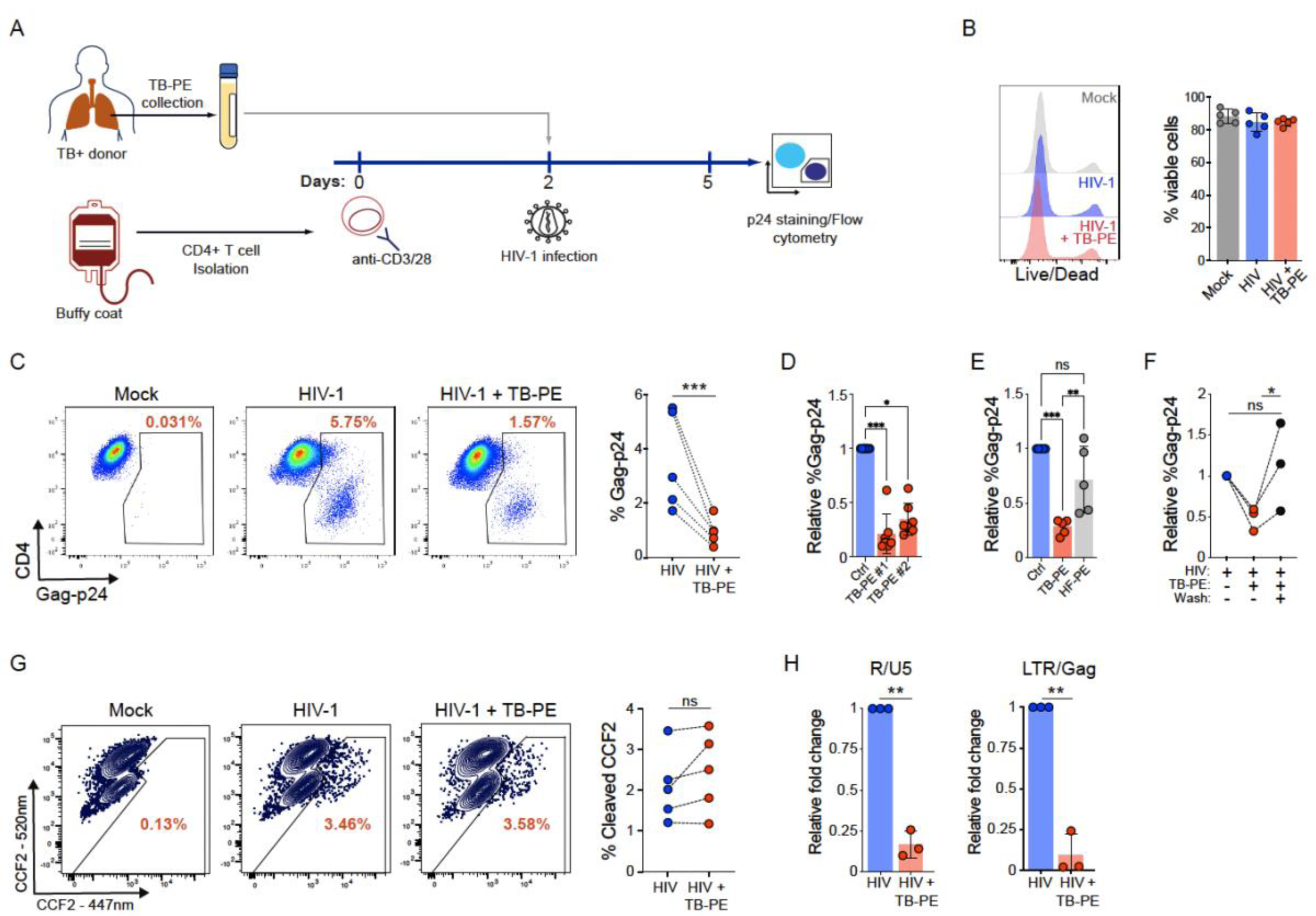
HIV-1 infection is inhibited by TB-PE in primary CD4+ T cells. Primary CD4+ T cells isolated from healthy donors were infected with HIV-1 in the presence of the acellular fraction of TB-PE. Cells infected with HIV-1 in the absence of TB-PE were used as a control. A) Schematic representation of the experimental design. B) Cell viability assessed by Live/Dead staining. C) The proportion of HIV-1 infected cells was determined at day 3 post-infection by Gag-p24 immunostaining. A representative cytometry (left) and Gag-p24 quantification (right) in cells from 5 independent donors are shown. D) Isolated CD4+ T cells from 7 healthy donors were infected with HIV-1 in the presence of TB-PE from 2 independent TB patients. The quantification of Gag-p24 positive cells, relative to HIV-1-infected cells in the absence of TB-PE, is shown. E) CD4+ T cells were treated with TB-PE or HF-PE and infected with HIV-1. The quantification of Gag-p24 positive cells relative to the control condition is shown. F) After 24 hours post-infection, a group of cells incubated with TB-PE were washed and cultured in fresh media without TB-PE. G) Viral entry was assessed by the HIV-1/BlaM-Vpr fusion assay. Enzymatic cleavage of CCF2 by BlaM-Vpr shifts the CCF2 fluorescence emission spectrum from 520nm to 447nm, indicating viral uptake. H) The HIV-1 reverse transcripts R/U5 and LTR/Gag were quantified by real-time qPCR at 6 h post-infection. Each dot represents values obtained from an independent donor. Statistical significance was determined by paired two-tailed t-test or One-way ANOVA followed by the Tukey’s HSD post-test. * p≤0.05, ** p≤0.01, ***p≤0.001.

### HIV-1 reverse transcription, but not viral entry, is inhibited by exposure of CD4+ T cells to TB pleural effusion

To identify which steps of the viral replication cycle were affected by a TB-associated microenvironment, we used the BlaM-Vpr fusion assay to assess viral entry in primary CD4+ T cells pretreated with TB-PE (33, 34). A shift in the CCF2 fluorescence emission spectrum from 520nm to 447nm indicates the enzymatic cleavage of CCF2 by Vpr-BlaM and therefore viral fusion with the plasma membrane of CD4+ T cells. Pre-incubation with TB-PE did not exhibit significant differences in the entry of HIV-1 when compared to the control condition (Figure 1G). Next, we quantified the levels of reversed-transcribed viral DNA to evaluate the impact of TB-PE on HIV-1 reverse transcription. Intriguingly, we observed that the reverse transcription of viral DNA was significantly decreased by TB-PE treatment (Figure 1H). These findings indicate that the microenvironment induced by *Mtb* infection impairs the replication of HIV-1 in CD4+ T cells by inhibiting viral reverse transcription.

### Transcriptional profile of CD4+ T cells exposed to TB pleural effusion

To understand the effect of the immune response to TB on CD4+ T cells and its potential impact on HIV-1 replication, we assessed the transcriptional profile of primary CD4+ T cells treated with TB-PE. Bulk RNA sequencing (RNAseq) was performed on primary CD4+ T cells isolated from 3 healthy donors and treated with TB-PE. In addition, since T cell activation increases CD4+ T cells susceptibility to HIV-1 infection (35), we also investigated the impact of TB-PE on CD4 T cells activated by anti-CD3/CD28 antibodies. As expected, the activation with anti-CD3/CD28 antibodies significantly changed the transcriptional profile of the stimulated CD4+ T cells as revealed by principal component analysis (PCA) (Figure 2A). Although PCA did not show consistent clustering patterns between untreated cells and cells incubated with TB-PE (Figure 2A), activated cells and TB-PE-treated activated cells clustered separately (Figure 2A). This result was further supported by the analysis of differentially expressed genes (DEGs) (Figure 2B-C; Supplementary Table 2). Moreover, Gene Set Enrichment Analysis (GSEA) revealed that cellular pathways involved in T cell activation were downmodulated by TB-PE treatment in anti-CD3/CD28-stimulated cells (Figure 2D-E, Supplementary Table 3).

**Figure 2.**
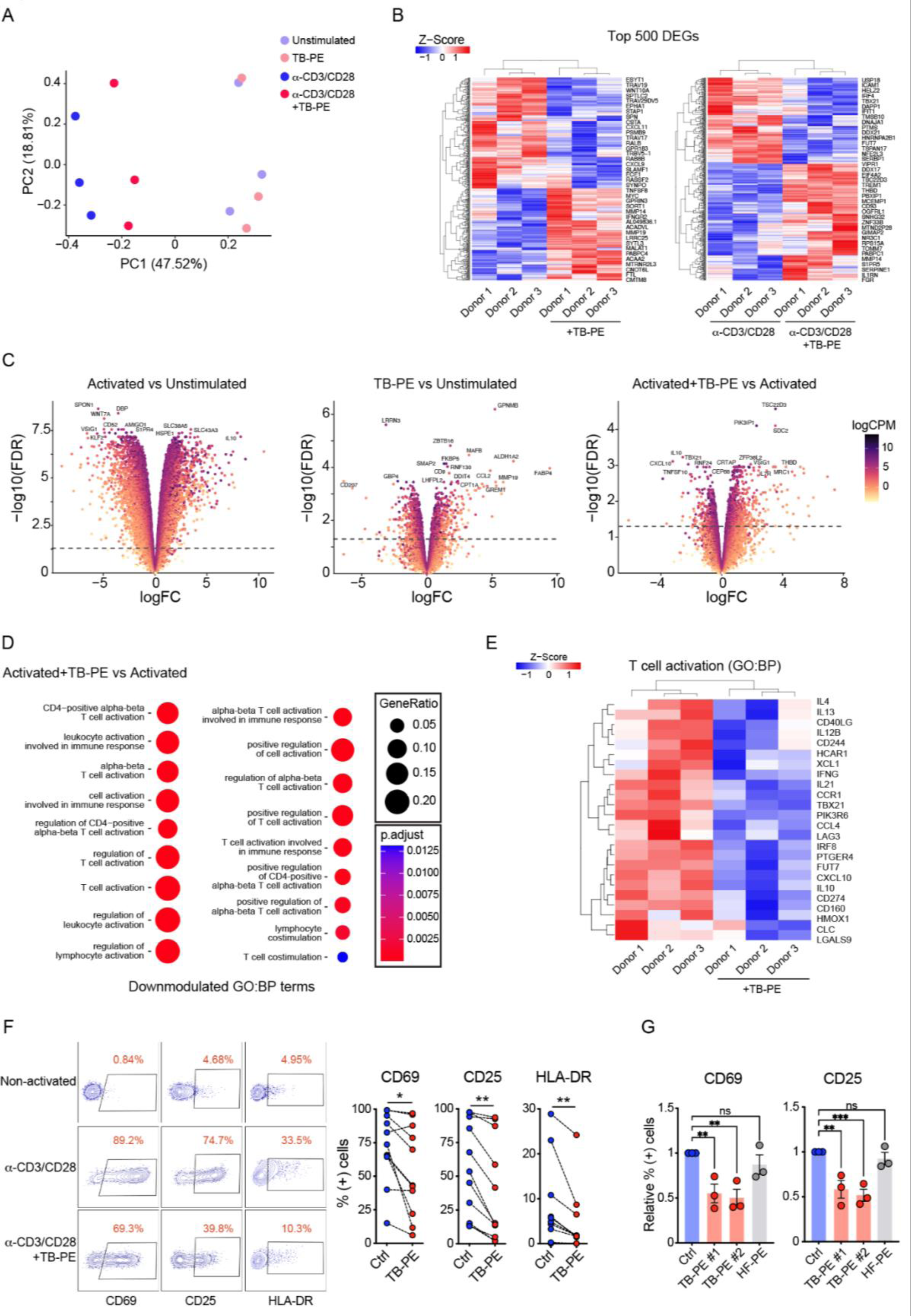
Transcriptional profile of CD4+ T cells exposed to TB-PE. The transcriptional profile of primary CD4+ T cells treated with TB-PE was characterized by bulk RNAseq analysis in resting and activated cells from 3 healthy donors. A) Principal component analysis of resting and activated CD4+ T cells. B) Heatmaps displaying hierarchical clustering of the top 500 variable genes between TB-PE-treated and control CD4+ T cells. Data from resting (left) and activated (right) CD4+ T cells are shown. C) Volcano plots representing the differentially expressed genes (DEGs) analysis. D) Gene set enrichment analysis (GSEA) comparing TB-PE-treated and control CD4+ T cells in resting and activated cells (DEGs: FDR<0.05 and absolute logFC>1). Dot plots illustrating the enriched GO biological processes terms related to T cell activation. The size of the dot reflects the gene ratio, indicating the proportion of differentially expressed genes associated with the pathway. The color intensity of the dot corresponds to the significance of enrichment, as determined by the adjusted p value. E) Heatmaps displaying the normalized expression of specific genes involved in T cell activation. Primary CD4+ T cells were stimulated with anti-CD3/CD28 antibodies in the presence of TB-PE or HF-PE. Cells activated in the absence of PEs were used as control. F) The expression of the activation markers CD69, CD25 and HLA-DR was measured by flow cytometry after 24 hours (CD69 and CD25) or 48 hours post-activation (HLA-DR). A representative cytometry (left) and the proportion of activated cells (right) are shown. G) Proportion of CD69 (left) or CD25 (right) positive cells stimulated in the presence of TB-PE from 2 independent TB-infected donors (red) or HF-PE (grey). Values are relative to the non-PE control condition (blue). Each dot represents values obtained from an independent donor. Statistical significance was determined by paired two-tailed t-test or One-way ANOVA followed by the Tukey’s HSD post-test. * p≤0.05, ** p≤0.01.

To validate our RNAseq results, we measured the expression of the activation markers CD69, CD25 and HLA-DR on primary CD4+ T cells upon the stimulation with anti-CD3/CD28 antibodies, in the presence or absence of TB-PE by flow cytometry. Consistent with our RNAseq analysis, TB-PE treatment resulted in a significant decrease in the surface expression of these activation markers (Figure 2F). A decrease in T cell activation was also observed when the CD4+ T cells were treated with individual TB-PE samples from two TB patients (Figure 2G). However, no significant inhibition of T cell activation was exhibited when these cells were incubated with HF-PE (Figure 2G). This result indicates that T cell activation can be downmodulated specifically by this TB-associated microenvironment. Our analysis shows that the TB-associated microenvironment, induced by the host anti-*Mtb* immune response, results in immunosuppressive properties, that lead to the downmodulation of the transcriptomic profile associated with T cell activation of CD4+ T cells.

### Oxidative phosphorylation and glycolysis are downregulated by TB pleural effusion treatment

Since T cell receptor (TCR)-mediated activation increases the levels of glycolysis and oxidative phosphorylation (OXPHOS) in CD4+ T cells, and as these two metabolic pathways are necessary for efficient HIV-1 replication (36–38), we also analyzed the metabolic profile of TB-PE-treated CD4+ T cells. We used the ATP rate assay and a Seahorse cell flux analyzer to measure oxygen consumption rate (OCR) and the proton efflux rate (PER) in CD4+ T cells as indicators of OXPHOS and glycolysis, respectively. The intracellular rate of ATP production derived from glycolysis (glycoATP) or OXPHOS (mitoATP) was also quantified. In line with the observed inhibition of T cell activation, both OXPHOS and glycolysis levels were significantly lower in TB-PE-treated cells when compared to controls (Figure 3A-B). Importantly, the incubation with HF-PE did not significantly impact either glycolysis or OXPHOS, indicating that this effect is specific to the TB-associated microenvironment.

**Figure 3.**
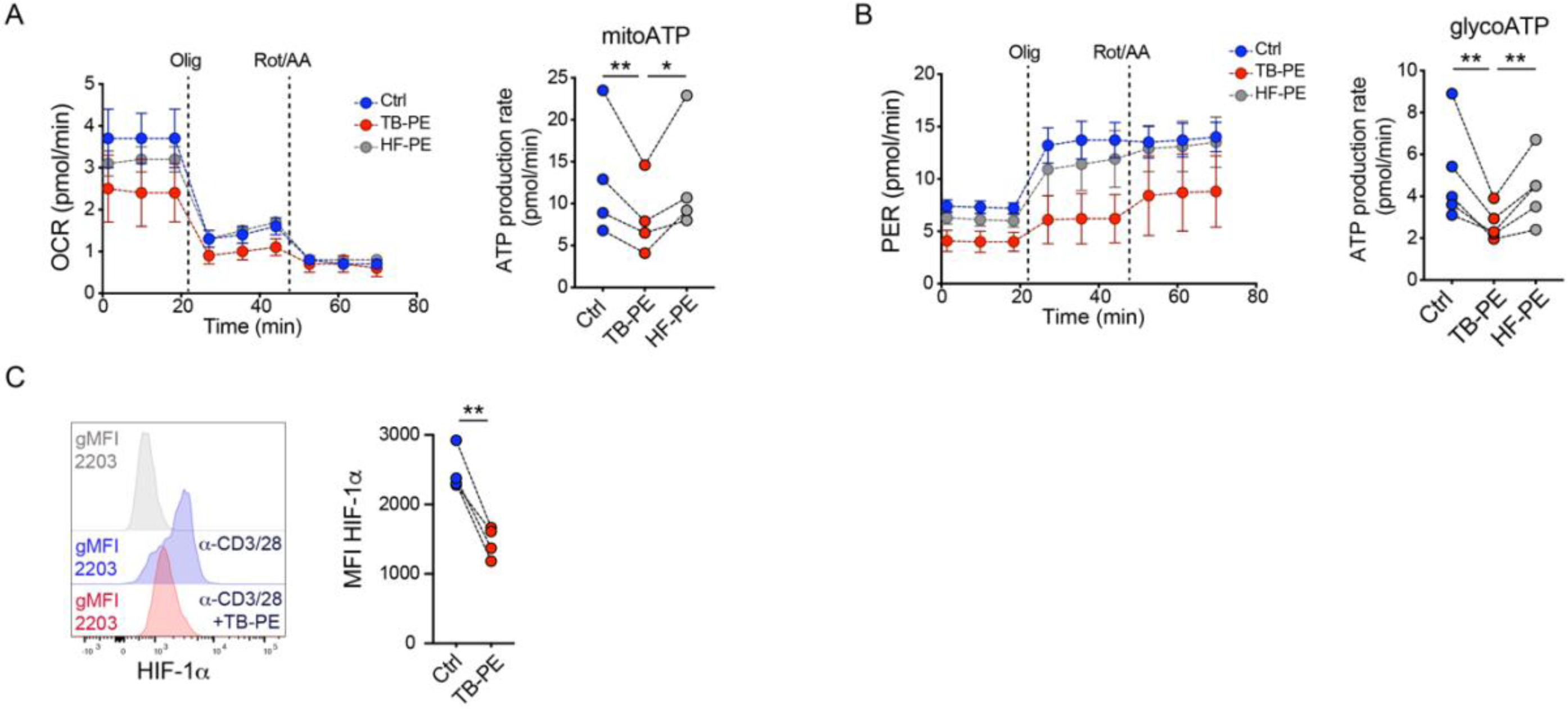
TB-PE decreases glycolysis and oxidative phosphorylation. Primary CD4+ T cells were stimulated with anti-CD3/CD28 antibodies in the presence of TB-PE or HF-PE. Cells activated in the absence of PEs were used as control. A) The expression of the activation markers CD69, CD25 and HLA-DR was measured by flow cytometry after 24 hours (CD69 and CD25) or 48 hours post-activation (HLA-DR). A representative cytometry (left) and the proportion of activated cells (right) are shown. B) Proportion of CD69 (left) or CD25 (right) positive cells stimulated in the presence of TB-PE from 2 independent TB-infected donors (red) or HF-PE (grey). Values are relative to the non-PE control condition (blue). C) Oxidative phosphorylation determined by Oxygen Consume Rate (OCR, left) and mitochondrial-dependent ATP production rate (right). D) Level of glycolysis was determined by Proton Efflux Rate (PER, left) and glycolysis-dependent ATP production rate (right). E) HIF-1α expression in activated CD4+ T cells was determined by flow cytometry. Each dot represents values obtained from an independent donor. Statistical significance was determined by paired two-tailed t-test or One-way ANOVA followed by the Tukey’s HSD post-test. * p≤0.05, ** p≤0.01.

Since we and others have shown that the Hypoxic Inducible Factor 1 alpha (HIF-1α) regulates and promotes glycolysis upon T cell activation and is also necessary for HIV-1 replication (39), we quantified the expression of HIF-1α in CD4+ T cells. Interestingly, the treatment with TB-PE significantly inhibited the expression of HIF-1α in activated CD4+ T cells (Figure 3C). These results suggest that the microenvironment found at the site of the TB infection inhibits metabolic pathways and factors necessary for efficient HIV-1 replication in CD4+ T cells.

### Cellular pathways involved in HIV-1 latency are modulated by TB pleural effusion treatment

The persistence of latently HIV-1 infected cells poses a significant barrier to eradicating the viral reservoir in PLWH. Therefore, we investigated the impact of the TB-associated microenvironment on HIV-1 latency in CD4+ T cells. Based on our RNAseq data obtained from activated CD4+ T cells, we analyzed cellular pathways necessary for efficient HIV-1 transcription and viral reactivation of latently HIV-1-infected cells (40–47). These cellular pathways include the nuclear factor kappa-light-chain-enhancer of activated B cells (NF-κB), Janus kinase/signal transducer and activator of transcription (JAK-STAT), tumor necrosis factor (TNF), extracellular signal-regulated kinases (ERK) and type I/II interferon (IFN) pathways(40–47). We found that NF-κB, JAK-STAT and type I/II IFN pathways were downmodulated in activated cells treated with TB-PE (Figure 4A-B). The NF-κB and type I/II IFN pathways were also downmodulated by TB-PE in non-activated CD4+ T cells (Supplementary Figure 1). Furthermore, we observed upregulation of pathways associated with transforming growth factor beta (TGFβ)-SMAD response, which is known to support HIV-1 latency (47–49), in the cells exposed to TB-PE (Figure 4 A-B). Of note, regulation of the ERK cascade signaling did not show consistent modulation patterns (Figure 4A). We also analyzed the expression of genes and pathways associated with programmed cell death, since previous studies indicate that intrinsic resistance to cell death of CD4+ T cells that compose the long-lived latent viral reservoir may contribute to HIV-1 persistence (47, 50–53). Our analysis revealed that programmed cell death pathways are inhibited by TB-PE (Supplementary Figure 2). This result suggests that CD4+ T cells exposed to TB-PE are less susceptible to cell death, which in turn could promote the persistence of latently HIV-1-infected cells.

**Figure 4.**
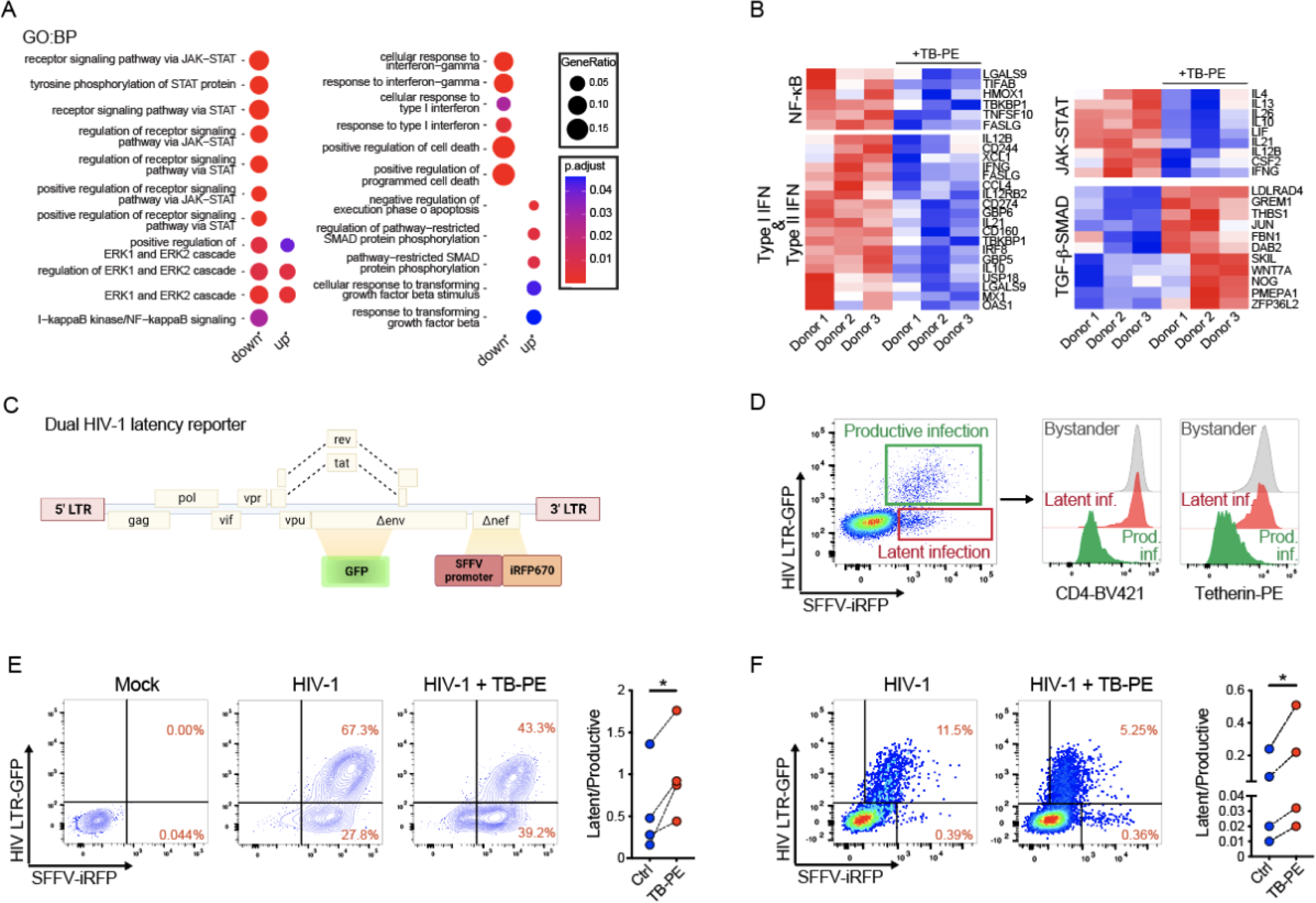
TB-PE promotes latency in CD4+ T cells. A) Gene set enrichment analysis (GSEA) comparing TB-PE-treated and control CD4+ T cells in activated cells (DEGs: FDR<0.05 and absolute logFC>1). The dot plots illustrate the positively and negatively enriched GO biological processes terms involved in HIV-1 transcription and reactivation. The size of the dot reflects the gene ratio, indicating the proportion of differentially expressed genes associated with the pathway. The color intensity of the dot corresponds to the significance of enrichment, as determined by the adjusted p value. B) Heatmaps displaying the normalized expression of specific genes involved in cellular pathways necessary for HIV-1 transcription and reactivation. C) Schematic of the dual HIV-1 latency reporter construct. D) CEM CD4+ T cells were infected with the dual HIV-1 latency reporter and the proportion productively (GFP+/iRFP+) and latent (iRFP+) infected cells was determined by flow cytometry (left). The expression of CD4 (middle) and Tetherin (right) on each gated population is shown. Jurkat (E) or primary CD4+ T cells (F) were infected with the dual HIV-1 latency reporter virus in the presence or absence of TB-PE. Representative cytometry (left) and the ratio between latent and productively infected cells (right) are shown. Each dot represents values obtained from an independent experiment (E) or independent donor (F). Statistical significance was determined by paired two-tailed t-test * p≤0.05.

### TB pleural effusion increases the number of latently HIV-1 infected CD4+ T cells

Several research groups have developed viral vectors containing the HIV-1 genome encoding a fluorescent reporter protein under the control of the viral LTR alongside a second reporter protein under the transcriptional control of an LTR-independent promotor (54–57). This strategy allows for the identification of latently and productively infected cells. Applying this strategy, we developed a dual HIV-1 latency reporter encoding two reporter proteins: the green fluorescent protein (GFP) and near-infrared fluorescent protein (iRFP) (Figure 4C). In our reporter HIV-1 construct, the expression of GFP is under the control of the viral LTR while iRFP expression is regulated by the spleen focus forming virus promoter (SFFV). We selected this promoter since SFFV is known to provide high-level transgene expression in primary human CD4+ T cells(58, 59). To test our new viral reporter of HIV-1 latency, we infected the CD4+ T cell line CEM, and measured the expression of GFP and iRFP by flow cytometry. The infection of CEM T cells with our dual reporter virus resulted in three populations: Double-positive for GFP and iRFP (productively infected cells), single-positive for iRFP (latently infected cells), and double-negative for both reporter proteins (uninfected bystander cells) (Figure 4D). To functionally validate our reporter virus, we assessed the expression of CD4 and Tetherin in the three populations. Since these two surface proteins are degraded by the viral protein Vpu, they should therefore be downmodulated in productively HIV-1-infected cells but not by cells latently infected with our reporter virus (60). As expected, GFP+iRFP+ cells efficiently downmodulated the expression of both CD4 and Tetherin (Figure 4D). However, iRFP+ cells expressed both markers at similar levels as bystander cells (Figure 4D) confirming a latent infection. After validating our reporter system of HIV-1 latency, we evaluated whether TB-PE increases the number of latently HIV-1 infected T cells. For this experiment, Jurkat CD4+ T cells were infected with our dual HIV-1 latency reporter in the presence or absence of TB-PE. Remarkably, the treatment with TB-PE increased the proportion of latently infected cells and reduced the proportion of productively infected cells (Figure 4 E). Importantly, this result was reproduced when primary CD4+ T cells were infected with our dual HIV-1 latency reporter (Figure 4F). Our findings indicate that the TB microenvironment promotes HIV-1 latency in CD4+ T cells.

### Reversal of HIV-1 latency is inhibited by TB pleural effusion treatment

Since our results showed that TB-PE promotes HIV-1 latency, we hypothesized that reactivation of latent HIV-1 may be impaired by this TB-associated microenvironment. To address our hypothesis, we used the J-Lat latency model. These Jurkat-derived cell lines are clones of cells containing one copy of integrated HIV-1 encoding the reporter GFP. Upon stimulation with latency reversing agents (LRAs) these cells express GFP, allowing for the quantification of HIV-1 reactivation by flow cytometry. To determine whether TB-PE affects the reactivation of latent HIV-1, we treated J-Lat cells (clones 6.3 and 10.6) with phorbol 12-myristate 13-acetate (PMA), a potent and well characterized LRA, in the presence or absence of TB-PE. In line with our previous results, the incubation with TB-PE inhibited the expression of GFP by both J-Lat clones (Figure 5A-B). The reactivation of HIV-1 was also inhibited when the cells were stimulated with latency reversal agents ingenol-3-angelate (PEP005) (both J-Lat clones; Figure 5A-B) and JQ1 (J-Lat clone 10.6; Figure 5B). Interestingly, TB-PE did not inhibit the reversal of HIV-1 latency induced by TNF-α (both J-Lat clones; Figure 5A-B) and Vorinostat (J-Lat clone 10.6; Figure 5B), indicating that TB-PE inhibition is pathway-specific. Of note, JQ1 and Vorinostat were used only on J-Lat clone 10.6 due to the cytotoxic effect of these LRAs on J-lat clone 6.3 (data not shown).

**Figure 5.**
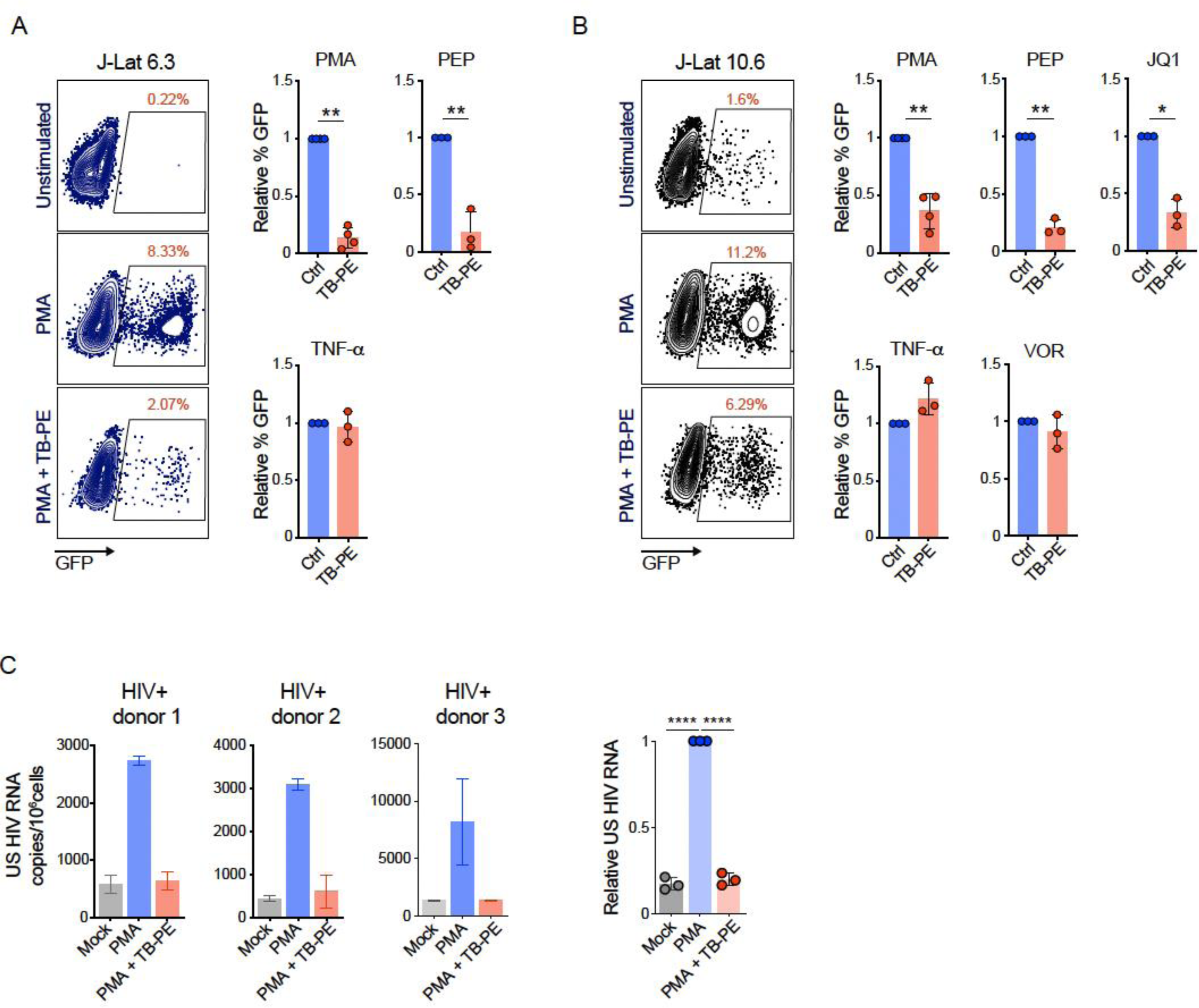
Reversal of HIV1 latency is impaired by TB-PE. J-Lat 6.3 (A) and 10.6 (B) cells were treated with different LRAs for 48 hours in the presence or absence of TB-PE and GFP expression was measured by flow cytometry as a correlate for viral reactivation. A representative cytometry (left) and GFP quantification relative to the control condition (right) are shown. A) J-Lat 6.3 cells were treated with 2ng/ml PMA, 50nM PEP005 or 10 ng/ml TNF-α. B) J-Lat 10.6 cells were treated with 0.5 ng/ml PMA, 0.05 ng/ml TNF-α, 5nM PEP005, 1μM JQ1 or 10μM Vorinostat. Each dot represents values obtained from an independent experiment. Statistical significance was determined by paired two-tailed t-test. C) Unspliced HIV-1 RNA (US HIV RNA) was measured in CD4+ T cells from 3 HIV-1-infected donors on ART after the stimulation with 2ng/ml PMA. Values obtained by real-time qPCR (left) and relative quantification of US HIV RNA copies per 10^6^ cells (right) are shown. Each dot represents values obtained from an independent donor. Statistical significance was determined by One-way ANOVA followed by the Tukey’s HSD post-test. * p≤0.05, ** p≤0.01, ****p≤0.0001.

Finally, to validate our *in vitro* results, we evaluated the effect of TB-PE *ex vivo* on cells obtained from PLWH. Isolated CD4+ T cells from 3 HIV-1-positive donors on ART were treated with PMA in the presence or absence of TB-PE. Levels of viral reactivation were quantified by measuring unspliced (US) HIV-1 RNA by quantitative PCR (qPCR). In agreement with our *in vitro* results, the incubation with TB-PE significatively inhibited viral reactivation of HIV-1 *ex vivo* (Figure 5C). Altogether, our results indicate that this TB-associated microenvironment induces viral latency and inhibits the reactivation of HIV-1 in latently infected CD4+ T cells.

## Discussion

While a growing body of evidence indicates worsened clinical outcomes in individuals coinfected with HIV-1 and *Mtb*, the precise interaction between these two pathogens remains unclear(61). The immunological interplay between HIV-1 and *Mtb* has predominantly focused on the virus-driven exacerbation of tuberculosis pathogenesis (62). However, our understanding of the mechanisms underlying the influence of pre-existing or subsequent *Mtb* infection on the replication, persistence, and progression of HIV-1, particularly in CD4+ T cells, is limited and requires further characterization. In this study, we investigated how a TB-associated microenvironment impacts HIV-1 infection and latency in CD4+ T cells. To further understand the interaction between both pathogens, in our study we use pleural effusion from TB patients, which reflects the microenvironment induced by *Mtb* infection and represents a relevant model for studying interactions at the site of the coinfection (21, 23, 32, 63).

Here, we show that TB-PE decreases HIV-1 replication in CD4+ T cells by inhibiting the reverse transcription of the viral genome, one of the early steps in the HIV-1 replication cycle. These results contrast with previous publications showing that TB-associated microenvironments or mycobacterial components increase HIV-1 replication and spread (19, 21, 24, 26, 64, 65). However, these studies have investigated HIV-1 replication in macrophages, indicating that the effect of *Mtb* infection and the TB-associated microenvironment varies between cell types (19, 21, 24, 26, 64, 65). Nevertheless, one study has shown that *Mtb* infection can also inhibit HIV-1 infection in monocyte-derived macrophages (25). Of note, He *et al.* have shown that blood-derived CD4+ T cells from individuals latently infected with *Mtb* supported HIV-1 replication more efficiently (66). Importantly, the authors did not detect this increase in viral infectivity when HIV-1 infection was assessed in cells from individuals undergoing active TB, which is when TB-PE is observed (66). The contrasting findings from these studies highlight the complexity of the HIV-1/TB interaction.

As mentioned above, the differences observed between our study and others can be explained in part by the cell type studied. In addition, our experimental model incorporates immune factors released in the microenvironment induced by *Mtb* infection. We observed that this microenvironment can modify the transcriptional profile of CD4+ T cells, particularly when these cells undergo TCR-dependent activation. T cell activation and downstream cellular events, such as increased rate of glycolysis and OXPHOS, as well as the expression of the metabolic regulator HIF-1α, were drastically inhibited by TB-PE. This inhibition is consistent with the observed decrease in HIV-1 infection since these cellular pathways and factors are required for efficient HIV-1 replication in CD4+ T cells (35–39, 67–70). In agreement with our results, it has been reported that immunosuppressive mediators present in pleural fluid from tuberculous pleurisy inhibit cytokine production, cell activation and Th1 differentiation of CD4+ T cells (71). Although we have focused our research on the impact of this immune microenvironment on HIV-1 replication and latency, our results suggest that diminished CD4+ T cell activation may also result in an impaired T cell-mediated immune response in coinfected individuals.

Even though a comprehensive understanding of viral latency is critical to progress towards an HIV-1 cure, the number of studies addressing this issue in the context of the HIV-1/TB coinfection remains limited to a few reports. For instance, recent studies have reported that components of the *Mtb* membrane and exosomes secreted by *Mtb*-infected macrophages reversed HIV-1 latency (72, 73). Moreover, it has also been shown that different mycobacterial components can modulate LTR-dependent transcription of HIV-1 in cell lines (18, 20), suggesting that these components may enhance HIV-1 transcription and reactivation. These reports contrast with our results showing that TB-PE promotes HIV-1 latency and impairs viral reactivation. However, as mentioned above, differences in our experimental model may explain these contrasting results. In line with the promotion of HIV-1 latency, our transcriptional profile analysis showed that cellular pathways necessary for HIV-1 transcription and reactivation, such as NF-κB and STAT (41, 42, 44), are downregulated by the TB-associated microenvironment. In addition, the TGF-SMAD pathway, reported to support HIV-1 latency (48, 49), was upregulated. Importantly, a decrease in T cell activation, OXPHOS and glycolysis has also been associated with the establishment of HIV-1 latency (47, 74–76), supporting our hypothesis. Interestingly, we observed that cell apoptosis is downmodulated in TB-PE-treated activated cells, indicating that this microenvironment may promote the survival of latently infected cells contributing to the persistence of HIV-1.

Since our results indicate that the immunosuppressive microenvironment generated by *Mtb* infection promotes HIV-1 latency, we hypothesize that the reactivation of HIV-1 might be impaired by this microenvironment. Indeed, we observed that TB-PE inhibits HIV-1 latency reversal in the cell line J-Lat and in cells from PLWH. However, this inhibition was restricted to some of the LRAs used in this study. For instance, HIV-1 reactivation by the PKC activators PMA (in both J-Lat cells and cells from PLWH) and PEP005 were drastically affected by TB-PE. This result is in line with our previous observations since PKC signaling is necessary for the TCR-triggered activation of T cells, which is dysregulated by TB-PE (77). In contrast, despite TB-PE treatment, HIV-1 latency was efficiently reversed by Vorinostat, a histone deacetylase (HDAC) inhibitor, and TNF-α. This is consistent with the finding that TB-PE mainly disrupts the TCR-mediated cell activation axes. However, JQ1, whose epigenetic mechanism of HIV-1 reactivation is by inhibiting the BET (bromodomain and extra-terminal domain) bromodomain BRD4 (bromodomain-containing protein 4), was also affected by TB-PE. This indicates that additional pathways can also be inhibited by this TB-associated microenvironment beyond those directly associated with T cell activation. This result indicates that some LRAs may be less effective in HIV-1/TB coinfected individuals.

Although our study provides mechanistic insights on the impact of the HIV-1/TB-coinfection on HIV-1 replication and latency, our findings rely on *in vitro* experimental data. However, by using TB-PE as well as CD4+ T cells obtained from people living with HIV-1, we have developed a physiologically-relevant model for researching the complex interactions between HIV-1 and the TB immune response. Therefore, the outcomes from this model contribute to relevant initial observations about HIV-1 persistence in CD4+ T cells in the context of HIV/TB coinfection. Another limitation is the use of lab-strain HIV-1 to study viral replication and latency. Hence, future research including primary strains of HIV-1 might be necessary to support the clinical relevance of our findings.

In conclusion, our results indicate that the immune response against *Mtb* infection leads to an anti-inflammatory microenvironment which promotes viral latency and persistence of HIV-1 at the site of the coinfection. While HIV-1 silencing mediated by this TB-associated microenvironment may reduce viral replication and spread at this specific anatomic site, its overall effect on the viral reservoir size may lead to negative clinical outcomes by stabilizing the HIV-1 reservoir and supporting its long-term persistence. Further research investigating the impact of the TB-associated microenvironment on the viral reservoir and anti-HIV-1 immune control is critical to complement and expand the findings of our study.

## Methods

### Study approval

This research was carried out in accordance with the Declaration of Helsinki (2013) of the World Medical Association. This study was approved by the institutional review boards at the Hospital F. J Muñiz (NIN-2601-19) and the Academia Nacional de Medicina de Buenos Aires (12554/17/X); Facultad de Medicina, Universidad de Buenos Aires (Buenos Aires, Argentina); and the Western Sydney Local Health District which includes the Westmead Institute for Medical Research (2022/PID00100 - 2022/ETH00092). Buffy coats from healthy donors were obtained from the Australian Red Cross Blood Service, Sydney, Australia. Written informed consent was obtained from all participants.

### Participant cohort and clinical samples

Pleural effusions were obtained by therapeutic thoracentesis at the Hospital F. J Muñiz (Buenos Aires, Argentina). The TB pleurisy diagnosis was based on a positive Ziehl– Neelsen stain or Lowestein–Jensen culture from pleural effusions and/or histopathology of pleural biopsy. The diagnosis was further confirmed by an *Mtb*-induced IFN-γ response and an adenosine deaminase-positive test (78). Exclusion criteria included a positive HIV-1 test, and the presence of concurrent infectious diseases or non-infectious conditions (cancer, diabetes, or steroid therapy). None of the patients had multidrug-resistant TB. Recruited patients who developed pleural effusion were divided in two groups according to their etiology (Supplementary Table 1). The first group included 9 patients diagnosed with tuberculous pleural effusion (TB-PE), while the second group comprised 4 patients with transudates resulting from heart failure (HF-PE) (Supplementary Table 1). For *ex vivo* HIV-1 reactivation assessment, individuals chronically infected with HIV-1 on ART were enrolled. The cohort of HIV-1-infected donors included for this study was defined as participants with established HIV-1 infection and on ART for at least 11 months (Supplementary Table 4).

### Pleural effusion samples

The pleural effusions were collected in heparin tubes and centrifuged at 300 g for 10 min at room temperature without brake. The cell-free supernatant was transferred into new plastic tubes, further centrifuged at 12,000 g for 10 min and aliquots were stored at −80°C. After diagnosis, pools were prepared by mixing equal amounts of individual PE derived from a specific etiology. The pools were decomplemented at 56°C for 30 min and filtered by 0.22 µm to remove remaining debris or residual bacteria. For the experimental conditions where pleural effusion was included, 20% v/v PE was added to the cell culture. This concentration of PE was determined as optimal by our group in previous studies (21, 23, 32, 63).

### Primary CD4+ T cells and cell lines

Peripheral blood mononuclear cells (PBMCs) were obtained by Ficoll-Hypaque density gradient centrifugation from buffy coats of healthy donors. CD4+ primary T cells were isolated and purified from PBMCs by negative selection using a CD4+ T cell isolation kit (MiltenyiBiotec). Cells were stimulated with anti-CD3/CD28 beads (MiltenyiBiotec) for two days in cRF10: RPMI (Lonza) supplemented with 10% fetal bovine serum (FBS, Gibco),1X GlutaMAX (Gibco), 100 U/ml penicillin (Gibco), 100 μg/ml streptomycin (Gibco), and 100 U/ml IL-2 (MiltenyiBiotec).

The J-Lat (clones 10.6 and 6.3) and Jurkat (E6-1) human CD4+ T cell lines were obtained from the NIH AIDS Reagent Program, Division of AIDS (NIAID, NIH). Cells were cultured in RF10: RPMI (Lonza) supplemented with 10% fetal bovine serum (FBS, Gibco),1X GlutaMAX (Gibco), 100 U/ml penicillin (Gibco) and 100 μg/ml streptomycin (Gibco). HEK 293T cells were obtained from the ATCC (CRL-11268) and cultured in DMEM (Lonza) supplemented with 10% fetal bovine serum (FBS, Gibco),1X GlutaMAX (Gibco), 100 U/ml penicillin (Gibco) and 100 μg/ml streptomycin (Gibco).

### Plasmids and HIV-1 viral strains

The molecular clones HIV-1 NL4-3AD8ENV (HIV^NL4-3^), pNL4-3-ΔEnv-EGFP and pHEF-VSVg were obtained through the NIH AIDS Reagent Program. The plasmids pCMV4-BlaM-Vpr, a gift from Warner Greene (Addgene plasmid # 21950), and pAdVAntage ™ Vector (Promega) were used for the HIV-1 fusion assay.

### Preparation of viral stocks

HIV^NL4-3^ and dual HIV-1 latency reporter viral stocks were produced by transfection of the corresponding vector (1 μg/well) in HEK 293T cells (3×10^5^ cells per well in 6-well plates) using X-treme gene transfection reagent (Roche). Pseudotyping was achieved by co-transfecting pHEF-VSVg (400 ng/well). HIV^NL4-3^/BlaM-Vpr viral stock was produced by co-transfecting HEK 293T cells (3×10^5^ cells per well in 6-well plates) with 1 μg NL4-3AD8ENV, 100 ng pCMV4-BlaM-Vpr and 60 ng pAdVAntage ™ Vector using X-treme gene transfection reagent (Roche). Supernatant was harvested at 48 and 72 h post-transfection, cleared by centrifugation at 1500 g for 10 min, and frozen at -80°C. The virus stocks were titrated by infecting Jurkat cells and flow cytometry analysis of p24 or GFP expression 72 h post-infection.

### HIV-1 infection

Isolated CD4+T cells (obtained as described above) were cultured at a concentration of 10^6^ cells/ml in cRF10 in a 96 well U-bottom plate and incubated overnight with the corresponding HIV-1 stock at 37°C. Then, cells were washed 2 times with cRF10, and fresh cRF10 was added. When indicated, the proportion of HIV-1-infected cells was determined by intracellular staining of the viral protein Gag-p24 with a Mouse anti-p24-FITC antibody (clone KC57, Beckman Coulter).

Cells infected with the dual HIV-1 latency reporter were spinoculated by centrifugation at 2200 rpm for 90 min in the presence of 8 μg/ml Polybrene and then incubated overnight at 37°C.

### Quantification of HIV-1 reverse transcripts

Total DNA was then extracted from HIV-1 infected cells at 6 h post-infection using the PureLink™ Genomic DNA Mini Kit (Thermo Fisher Scientific). Quantitative PCR for R-U5 and LTR-Gag transcripts was performed using the following primers:

R/U5 Fwd: 5’-GGCTAACTAGGGAACCCACTG-3’

R/U5 Rev: 5’-CTGCTAGAGATTTTCCACACTGAC-3’

LTR/Gag Fwd: 5’-TGTGTGCCCGTCTGTTGTGT-3’

LTR/Gag Rev: 5’-GAGTCCTGCGTCGAGAGAGC-3’

β2m Fwd: 5’-TGCTGTCTCCATGTTTGATGTATCT-3’

β2m Rev: 5’-TCTCTGCTCCCCACCTCTAAGT-3’

Quantitative PCR (qPCR) was performed using the Quantitect SYBR green PCR Master mix (Roche) with 10 ng of DNA and 0.5 μM primers in a final volume of 10 μl. DNA amplification was performed using the CFX96 Touch Real-Time PCR Detection System (Bio-Rad). Each sample was amplified in triplicate and relative expression was calculated by normalization to β2m.

### HIV-1 fusion assay

HIV-1 fusion assay was adapted from Cavrois et al. (33). Briefly, cells were incubated 4 h with HIV^NL4-3^/BlaM-Vpr at 37°C. After incubation, cells were washed with CO_2_ independent medium (Gibco) and incubated 1 hour with 100 μl of CCF2-AM loading solution (Thermo Fisher Scientific) at room temperature in the dark. Next, cells were washed once with CO_2_ independent media and incubated overnight in 200 μl of development media (Thermo Fisher Scientific) at room temperature. CCF2-AM loading solution and development media were prepared as previously indicated (34). After the overnight incubation, cells were washed 3 times with PBS and fixed in 1% paraformaldehyde (PFA) for 20 min at 4°C. After fixation, the cells were washed with PBS twice, resuspended in 200 μl PBS and acquired on a LSRFortessa Flow Cytometer. Enzymatic cleavage of CCF2 by Vpr-BlaM shifts the CCF2 fluorescence emission spectrum from 520nm to 447nm, indicating viral uptake.

### Dual HIV-1 latency reporter

To generate the dual HIV-1 latency reporter vector, pNL4-3-ΔEnv-EGFP was digested with HpaI and XhoI. Then, a synthetic gene fragment containing the Spleen-forming focus virus promoter sequence followed by the iRFP670 protein-coding sequence (gBlock; Integrated DNA Technologies, IDT) was assembled with the gel-purified plasmid using the HiFi Assembly Master Mix (NEB, E5520) disrupting *nef* sequence (Δnef). The vector sequence was verified by Sanger sequencing (Source BioScience).

### RNAseq library

Primary CD4+ T cells were obtained as described above. A group of isolated CD4+ T cells were activated using anti-CD3/CD28 antibodies in the presence or absence of TB-PE, and after 20 hours, the cells were harvested to perform bulk RNAseq. RNA from CD4+ T cells (2×10^6^ cells per condition) was extracted by using the RNeasy Plus Mini kit (QIAGEN). Extracted total RNA was quantified using the Qubit broad range RNA assay, and RNA quality was assessed using the Agilent 4200 Tapestation system. Library preparation was performed using 400ng of total RNA as input and the Illumina stranded mRNA Prep kit. Short read sequencing was performed on the Novaseq 6000 platform where a minimum of 20 million paired-end 50bp reads per sample were generated.

### RNAseq analysis

Library sequencing quality was determined using FastQC (Babraham Bioinformatics: www.bioinformatics.babraham.ac.uk/). Illumina adaptor sequence and low quality read trimming (read pair removed if < 20 base pairs) was performed using Trim Galore (BabrahamBioinformatics). STAR (79) was used to align reads to human genome hg38 using ENSEMBL gene annotations as a guide. Read counts data corresponding to ENSEMBL gene annotations were generated the STAR flag --quantMode GeneCounts. Multiqc(80) was used to verify quality metrics. All analyses were performed in the R Statistical Environment ((81); http://www.Rproject.org) with the Tidyverse package (82). EdgeR was used to perform background correction and normalize count data by library size (83). Differentially expressed gene analysis was determined using the QLFtest (84) (BH MTC p<0.05). Gene lists were functionally annotated with Gene Ontology Biological Process (GO:BP) pathways (adjusted p value<0.05) using the clusterProfiler package(85).

### *In vitro* reactivation of HIV-1

J-Lat cells, clones 10.6 and 6.3, were plated at a concentration of 3×10^5^ cell per well in a 96-well plate and stimulated with the indicated latency reversal agent (LRA) for 48h. Every LRA was titrated individually in both J-Lat clones to determine the optimal experimental concentration. To reverse HIV-1 latency in J-Lat cells clone 10.6, 0.5 ng/ml PMA, 0.05 ng/ml TNF-α, 5nM PEP005, 1μM JQ1 and 10μM Vorinostat were used. To reverse HIV-1 latency in J-Lat cells clone 6.3, 2 ng/ml PMA, 10 ng/ml TNF-α and 50nM PEP005 were used. JQ1 and Vorinostat were not used in J-Lat cells clone 6.3 due to the high cytotoxicity observed in this cell clone when these LRAs were titrated. Cell viability was determined by Live/Dead near-infrared (Invitrogen) staining and HIV-1 reactivation by quantifying GFP expression by flow cytometry.

### *Ex vivo* reactivation of HIV-1

*Ex vivo* HIV-1 latency reversal was assessed as previously described (86). Briefly, CD4+ T cells were isolated from 40 ml of peripheral blood from HIV-1-infected individuals on ART by immunodensity negative-selection using the RosetteSep™ Human CD4+ T Cell Enrichment Cocktail (STEMCELL Technologies). Cells were plated at a concentration of 1-3×10^6^/ml in RF10 and stimulated with 2 ng/ml PMA for 16 h when indicated. RNA was extracted by using the PureLink RNA minikit (Invitrogen) and reverse transcribed using the SuperScript™ III Reverse Transcriptase (Invitrogen) following the manufacturer’s recommendations. HIV-1 unspliced RNA (US-RNA) quantification as previously described (87). Briefly, a heminested PCR was run for 15 cycles and a second amplification round was performed by quantitative real-time PCR for 40 cycles. US-RNA copy number was normalized to 18S RNA expression. Each sample was assessed in triplicate, and a non-reverse-transcribed control was used to detect DNA contamination.

### Glycolysis and oxidative phosphorylation rate measurement

CD4+ T cells were purified by magnetic isolation and stimulated with anti-CD3/CD28 antibodies in the presence or absence of TB-PE. After 24 h, the cells were cultured in RPMI-based Seahorse medium (Agilent) supplemented with 10 mM D-glucose, 1 mM pyruvate and 2 mM glutamine and plated in a Seahorse 24-well plate at a concentration of 7×10^5^ cells/well. Proton efflux rate (PER) and oxygen consumption rate (OCR) were measured using a seahorse XFe24 analyzer. ATP production rates were measured using the Seahorse XF Real-Time ATP Rate Assay kit. Oligomycin, rotenone and antimycin A were added to quantify mitochondrial-derived and glycolysis-derived ATP production rate. OCR and PER values were normalized by calculating the cellular area per condition.

### Statistics

The experimental data was analyzed using Prism (GraphPad Software). Normality of the data was tested using the Kolmogorov–Smirnov test. Based on the normality test, either one-way ANOVA followed by the Tukey’s HSD post-test or Kruskal–Wallis followed by Dunn’s post-test were used for multiple comparison analyses and a Student’s t-test used to compare two conditions. A p-value less than 0.05 was considered significant.

## Acknowledgments

We acknowledge with gratitude the participants who donated samples for this study. We thank Drs. Natalia Laufer, Xavier Aragone and Milagro Sanchez Cunto for enrolling the participants for this study. Flow cytometry and RNAseq analysis were performed at the Westmead Scientific Platforms, which is supported by the Westmead Research Hub, the Cancer Institute New South Wales, the National Health and Medical Research Council, and the Ian Potter Foundation. We would like to thank Dr. Joey Lai, Genomics Facility manager at The Westmead Institute for Medical Research, for the RNA-seq library preparation. This work was supported by the Delaney AIDS Research Enterprise to Find a Cure (1UM1AI126611-01 and 1UM1AI164560-01), the Australian National Health and Medical Research Council (APP1149990), the *Agence Nationale de Recherche sur le Sida et les hépatites virales*(ANRS2018-02, ECTZ 118551/118554, ECTZ 205320/305352, ANRS ECTZ103104 and ECTZ101971), *Sidaction* (13457), the Argentinean National Agency of Promotion of Science and Technology (PICT 2019-2019-01044), Sandra and David Ansley, the Sydney Medical School Foundation, and The University of Sydney Institute for Infectious Diseases.

## Author contributions

G.D., L.B. and S.P. conceptualized and designed the study. S.C., A.V. and G.D. designed and conducted the majority of ex vivo and in vitro experiments. A.C. and G.T. evaluated ex vivo HIV reactivation. C.M. and P.P.G. developed the Dual HIV-1 Latency Reporter. B.G. and T.O. contributed to the transcriptomic analysis. M.S. assisted with in vitro experiments. Z.V. and C.V. performed metabolic assays. G.D. wrote the original manuscript. S.C., L.B. and S.P. supervised and edited the original manuscript. Co–first authorship order was determined by the contribution to the results presented in the manuscript and contribution to writing the initial manuscript draft.

